# Early life stress dysregulates kappa opioid receptor signaling within the lateral habenula

**DOI:** 10.1101/2020.07.15.202614

**Authors:** Sarah C. Simmons, Ryan D. Shepard, Shawn Gouty, Ludovic D. Langlois, Brian M. Cox, Fereshteh S. Nugent

**Affiliations:** Uniformed Services University of the Health Sciences, Edward Hebert School of Medicine, Department of Pharmacology and Molecular Therapeutics, Bethesda, Maryland 20814, USA

**Author notes:** **Corresponding author:** Fereshteh S. Nugent.

**Keywords:** lateral habenula, LHb, kappa opioid receptor, KOR, dynorphin, early life stress

## Abstract

The lateral habenula (LHb) is an epithalamic brain region associated with value-based decision making and stress evasion through its modulation of dopamine (DA)-mediated reward circuitry. Specifically, increased activity of the LHb is associated with drug addiction, schizophrenia and stress-related disorders such as depression, anxiety and posttraumatic stress disorder. Dynorphin (Dyn)/Kappa opioid receptor (KOR) signaling is a mediator of stress response in reward circuitry. Previously, we have shown that maternal deprivation (MD), a severe early life stress, increases LHb intrinsic excitability while blunting the response of LHb neurons to extra hypothalamic corticotropin-releasing factor (CRF) signaling, another stress mediator. CRF pathways also interact with Dyn/KOR signaling. Surprisingly, there has been little study of direct KOR regulation of the LHb despite its distinct role in stress, reward and aversion processing. To test the functional role of Dyn-KOR signaling in the LHb, we utilized ex-vivo electrophysiology combined with pharmacological tools in rat LHb slices. We show that activation of KORs by a KOR agonist (U50,488) exerts differential effects on the excitability of two distinct subpopulations of LHb neurons that differ in their expression of hyperpolarization-activated cation currents (HCN, Ih). Specifically, KOR stimulation increases neuronal excitability in LHb neurons with large Ih currents (Ih+) while decreases neuronal excitability in small/negative Ih (Ih-) neurons. Additionally, we found that an intact fast-synaptic transmission is required for the effects of U50,488 on the excitability of both Ih- and Ih+ LHb neuronal subpopulations. Consistently, KOR activation also altered both glutamatergic and GABAergic synaptic transmission. While stimulation of presynaptic KORs uniformly suppressed glutamate release onto LHb neurons, we found that U50, 488 either increased or decreased GABA release. We also found that MD significantly increased immunolabeled Dyn (the endogenous KOR agonist) labeling in neuronal fibers in LHb while significantly decreased mRNA levels of KORs in LHb tissues compared to those from non-maternally deprived (non-MD) control rats. While total p38 MAPK (a downstream signaling pathway driven by KOR activation) expression was elevated in the LHb of MD rats compared to non-MD controls, we found that application of KOR-specific agonist, U50,488, onto LHb slices was still able to alter phosphorylated p38 MAPK (ph-p38) expression in MD rats similar to non-MD controls. Moreover, we found that the U50,488-mediated increase in LHb neuronal firing observed in non-MD rats was absent following MD. Altogether, this is the first demonstration of the existence of the functional Dyn/KOR signaling in the LHb that can be modulated in response to severe early life stressors such as MD.

## 1. Introduction

Depression is a major health concern in the United States with a prevalence of 7% of all adults or 13% of all adolescents (age12-17) in 2017 [1, 2]. Those who experience severe early life stress (ELS) or adverse childhood events (ACE) have increased risk for the development of depression, suicide ideation and suicide attempts later in life [3–6]. 61% of the US population have experienced at least 1 ACE, and16% have experienced 4 or more, putting a significant population at increased risk for developing mood-disorders later in life [3]. In order to better understand the interaction of ELS/ACE and increased susceptibility to development of a mood-disorder (such as anxiety or depression) we use a preclinical rodent model of severe childhood neglect – maternal deprivation (MD). In this model, neonatal rat pups are separated from the dam for a single 24hr period which induces long-lasting effects on depressive-like behavior that can be reversed by common antidepressant treatments effective in treating depression in human subjects [7–11]. Additionally, we have found that MD alters the activity of key reward/aversion processing regions such as the ventral tegmental area (VTA) and the lateral habenula (LHb) [7–9, 12, 13]. The LHb is an epithalamic brain region that is now recognized as an emerging anti-reward hub implicated in motivation and decision-making. Neuronal pathways through LHb link forebrain limbic structures with midbrain monoaminergic centers [14–18]. LHb is involved in reward/aversion-related learning and memory processes associated with avoidance from stressful and aversive situations through suppression of dopamine (DA) and serotonin (5HT) systems and its increased activity is associated with depression in human studies and depression-like phenotypes across a variety of animal models of stress [7, 13, 19–24].

The dynorphin-Kappa opioid receptor (Dyn/KOR) system is highly implicated in regulating states of motivation and emotion, and in the development of mood disorders, including depression and anxiety, as well as in drug addiction and stress-induced relapse to drug-taking [25–30]. Activation of Dyn/KOR signaling in response to acute stress increases motivation to escape an immediate threat, whereas prolonged activation of this signaling during chronic stress leads to dysregulation of this neuromodulatory system that could also act as an anti-reward system by producing aversion that counteracts positive reinforcement and contributes to the development of pro-depressive, anti-social and negative affective and emotional states associated with mood/stress-related disorders as well as addiction [27]. While there is evidence for direct Dyn/KOR regulation of the mesocorticolimbic system [25, 31–33], to date there has been little study of KOR regulation of other important stress/mood-related regions such as the LHb. Dyn/KORs are abundant in reward and aversion brain structures projecting to the LHb. For example, the LHb receives significant inputs from pro-dynorphin expressing brain regions with high levels of KOR expression such as lateral hypothalamus, amygdala, and nucleus accumbens (NAc), dorsal striatum, and globus pallidus internal (also referred to as the entopeduncular nucleus in rodents) [23, 26, 34–38]. KOR expression has also been reported in the LHb in addition to mu opioid receptors (MOR) and delta opioid receptors (DOR)[39, 40], also raising the possibility of postsynaptic modulation of LHb neurons by Dyn/KOR signaling in addition to presynaptic effects through inputs projecting to the LHb. Therefore, the LHb is poised to be not only a potent regulator of affective disorders and addiction, but also a possible hotspot for Dyn/KOR regulation of motivation, stress and mood.

Here we first established the role of Dyn/KOR signaling in modulating LHb activity through both intrinsic properties and synaptic modulation. We provide significant evidence for the presence of a functional Dyn/KOR signaling in the LHb that individually alters the activity of two discrete populations of LHb neurons discriminated by the presence or absence of hyperpolarization-activated cation currents (HCN, Ih). Additionally, we found that KOR activation in LHb directly modulated glutamatergic and GABAergic synaptic transmission in the LHb, contributing to changes in overall LHb excitability. Secondly, we assessed whether this Dyn/KOR regulation of LHb neuronal excitability was altered following MD. We found that MD dysregulated Dyn/KOR signaling, resulting in a loss of acute KOR modulation of LHb neuronal excitability. To our knowledge, this is the first study directly assessing the effects of Dyn/KOR signaling in the LHb, and providing evidence for Dyn/KOR dysregulation in the LHb following a severe ELS.

## 2. Methods

### 2.1. Animals

All experiments employed male Sprague Dawley rats (sourced from Taconics Inc, and Charles River) in experiments conducted in accordance with the National Institutes of Health Guide for the Care and Use of Laboratory Animals and were approved by the Uniformed Services University Institutional Animal Care and Use Committee. All rats were received on post-natal day 6 (PN6) with lactating dam and allowed to acclimate undisturbed to a reverse 12hr dark: 12hr light cycle schedule with lights on at 18:00 for 48hrs, before initiation of the maternal deprivation (MD) procedure. All procedures began 3-4 hours after the start of the light-cycle, and all animals received ad-lib standard chow and water (except where noted during the MD procedure). All efforts were made to minimize animal suffering and reduce the number of animals used throughout this study.

### 2.2. MD procedure

MD was performed on male rats at PN9. Half of a litter (randomly selected) were isolated from the dam and their siblings for 24hrs (MD group). The isolated pups were placed on a heating pad (34°C) in a separate quiet room and not disturbed for 24hr until being returned to their home cage. The remaining non-maternally deprived control group (non-MD) received the same amount of handling. For electrophysiology, western blot, and qPCR experiments all rat pups were sacrificed during the adolescent age range of PN21-PN28. For IHC experiments, rats were sacrificed at ages PN16 (young), PN28 (adolescent), and PN60 (adult). For IHC of adult animals, all rats were co-housed (2 per cage, treatment matched) from weaning at PN28 to sacrifice at PN60 and received standard housing care but no additional experimenter manipulation prior to sacrifice.

### 2.3. Slice Preparation

For all electrophysiology experiments several separate cohorts of non-MD/MD treated rats were used. As described before [13] all rats were anesthetized with isoflurane, decapitated and brains were quickly dissected and placed into ice-cold artificial cerebrospinal fluid (ACSF) containing (in mM): 126 NaCl, 21.4 NaHCO_3_, 2.5 KCl, 1.2 NaH_2_PO_4_, 2.4 CaCl_2_, 1.00 MgSO_4_, 11.1 glucose, 0.4mM ascorbic acid and saturated with 95% O_2_-5% CO_2_. Briefly, sagittal slices containing LHb were cut at 250μm and incubated in above prepared ACSF at 34°C for at least 1 hour prior to electrophysiological experiments. For patch clamp recordings, slices were then transferred to a recording chamber and perfused with ascorbic-acid free ACSF at 28°C.

### 2.4. Electrophysiology

Voltage-clamp cell-attached and whole-cell voltage/current-clamp recordings were performed from LHb neurons in LHb-containing slices using patch pipettes (3-6 MOhms) and a patch amplifier (MultiClamp 700B) under infrared-differential interference contrast microscopy. Data acquisition and analysis were carried out using DigiData 1440A, pCLAMP 10 (Molecular Devices), Clampfit, and Mini Analysis 6.0.3 (Synaptosoft Inc.). Signals were filtered at 3 kHz and digitized at 10 kHz.

To assess LHb spontaneous activity, LHb neuronal activity/excitability and intrinsic membrane properties in intact synaptic transmission, cells were patch clamped with potassium-gluconate based internal solution (130 mM K-gluconate, 15 mM KCl, 4 mM adenosine triphosphate (ATP)-Na+, 0.3 mM guanosine triphosphate (GTP)-Na+, 1 mM EGTA, and 5 mM Hepes, pH adjusted to 7.28 with KOH, osmolarity adjusted to 275 to 280 mOsm) in slices perfused with ACSF. In a separate set of experiments, intrinsic neuronal excitability experiments were performed in the absence of fast-synaptic transmission by addition of AMPAR antagonist 6,7 dinitroquinoxaline-2,3-dione di-sodium salt (DNQX; 10μM,Tocris-2312), GABAAR antagonist picrotoxin (100 μM, Tocris-1128), and NMDAR antagonist D-(-)-2-Amino-5-phosphonopentanoic acid 50 μM, Tocris-0106) to the ACSF. Spontaneous neuronal activity and AP firing patterns (tonic, bursting) were assessed in both cell-attached recordings in voltage-clamp mode at V=0mV and whole cell recording in current-clamp mode at I=0 pA for the duration of ~1min recording. Cells that fired less than 2 action potentials (AP) during this time period were characterized as silent. Resting membrane potential (RMP) was assessed immediately after achieving whole-cell patch configuration in current clamp recordings. Ih current recordings were performed in voltage-clamp in response to stepping cells from −50 mV to −100 mV (700ms duration). Ih positive (Ih+) neurons were defined as having a maximal Ih current (peak to steady state trough) response > – 10pA (and conversely Ih-neurons as ≤ −10pA).

During LHb neuronal excitability recordings in current-clamp mode, AP generation was assessed in response to increasingly depolarizing current steps ranging from +10 to +100pA (+10 pA ea. step) while cells were kept at −67 to −70mV with manual direct current injection between pulses. Current steps were 5s in duration with 25s inter-stimulus intervals. The number of APs induced by depolarization at each intensity was counted and averaged for each experimental group at each step. Input resistance (Rin) was measured at −50pA step (5s duration) and calculated by dividing the steady-state voltage response by the current-pulse amplitude and presented as MOhms (MΩ). AP number, AP threshold, fast afterhyperpolarizations (fAHP), medium after-hyperpolarizations (mAHP), AP half-width, AP amplitude were assessed using clampfit and measured from the first AP at the current step that was sufficient to generate the first AP/s as previously described [11].

Whole cell recordings of AMPAR-meditated miniature excitatory postsynaptic currents (mEPSC) were performed in ACSF perfused with picrotoxin (100 μM), d-APV (50μM), and tetrodotoxin (TTX, 1 μM). Patch pipettes for mEPSCs recordings were filled with cesium (Cs)-gluconate internal solution (117mM Cs-gluconate, 2.8 mM NaCl, 5mM MgCl_2_, 2mM ATP-Na+, 0.3 mM GTP-Na+, 0.6 mM EGTA, and 20mM Hepes, pH adjusted to 7.28 using CsOH, osmolarity adjusted to 275-280 mOsm). Whole-cell recordings of GABA_A_R-mediated miniature inhibitory postsynaptic currents (mIPSC) were performed in ACSF perfused with AP-V (50uM), DNQX (10 μM), glycine receptor inhibitor (strychnine, 1 μM), and TTX, 1 μM. Patch pipettes were filled with KCl internal solution (125 mM KCl, 2.8 mM NaCl, 2 mM MgCl_2_, 2mM ATP Na+, 0.3 mM GTP-Na+, 0.6 mM EGTA, and 10mM Hepes, pH adjusted to 7.28 with KOH, osmolarity adjusted to 275-280mOsm). For both mEPSCs and mIPSCs, LHb neurons were voltage-clamped at −70mV and recorded over 10 sweeps, each lasting 50s. The cell series resistance was monitored through all the experiments and if this value changed by more than 10%, data were not included.

### 2.5. Drugs

For all agonist/antagonist experiments a within-subjects experimental design was used. Baseline recordings were first performed (depolarization-induced AP recordings /mEPSC /mIPSC) for each neuron and then appropriate drug was added to the slice by the perfusate and responses tested following 30-45min of drug bath application. In all experiments stimulating KORs, the KOR agonist (+/-)U50,488 (Tocris, 10 μM) was used. To test specificity of U50,488 effects, we used a novel selective KOR antagonist (BTRX-084266, 10-20μM) generously provided by BlackThorn, Inc. LHb-containing slices were incubated with BTRX-084266 for at least 30min prior to U50,488 application.

### 2.6. Immunohistochemistry and image analyses

Rats were anesthetized with an intraperitoneal injection containing ketamine (85 mg/kg) and xylazine (10 mg/kg) and perfused through the aorta with 1x phosphate buffered saline (PBS), followed by 4% paraformaldehyde (PFA) (Santa Cruz). Juvenile, adolescent or adult rats (PN16, PN21-28 or PN60) were perfused similarly but with adjusted volumes of between 200-250ml each solution. The brains were dissected and placed in 4% PFA for 24 h and then cryoprotected by submersion in 20% sucrose for 3 days, frozen on dry ice, and stored at −70°C until sectioned. Sections of the LHb were cut using a cryostat (Leica CM1900) and mounted on slides. Serial coronal sections (20 mμ) of the midbrain containing the LHb (from −2.64 to −4.36 mm caudal to bregma (Paxinos and Watson, 2007) were briefly fixed in 4% PFA for 5 min, washed in 1x PBS, and then blocked in 10% normal goal serum (NGS) containing 0.3% Triton X-100 in 1x PBS for 1 h. Sections were incubated in rabbit anti-Dynorphin A 1-8 (Dyn-A, 1:500,[37, 41–44]) and guinea pig anti-NeuN (1:500, Synaptic Systems) in carrier solution (5 % NGS in 0.1% Triton X-100 in 1x PBS) overnight at room temperature. After rinsing in 1x PBS, sections were incubated for 2 h in Alexa Flour 488-labeled goat anti-rabbit IgG and Alexa Flour 647-labeled goat anti-guinea pig IgG (both diluted 1:200). Finally, sections were rinsed in 1x PBS, dried, and coverslipped with Prolong^®^ mounting medium containing DAPI to permit visualization of nuclei. Background staining was assessed by omission of primary antibody in the immunolabeling procedure (negative control). LHb tissue sections of rats with previously established presence of Dyn-A / NeuN immunoreactive neurons were processed as positive control tissue. Images were captured using a Leica DMRXA Florescent Microscope.

Two 40x images of the LHb were taken at three AP location (−3.8, −3.9, and −4.1 relative to bregma). Using ImageJ sample pixel intensity, for each image, was averaged to include all pixels above of what was considered the “true” signal (35 pixel intensity). Background pixel averaging was performed to include all pixels below 35 pixel intensity. Density readings were created by normalizing all sample readings to background. The two normalized density reading from each AP location were averaged, and then each animal was averaged across AP location before group means were calculated.

### 2.7. Western Blot and qPCR

For all western blot experiments we used similar procedures as published previously [11–13]. Specifically, adolescent rats (PN21-28) were anesthetized using isofluorane and immediately decapitated. Brains were quickly dissected and placed in ice-cold ASCF and sagittal slices were cut at 300μm using a vibratome. All control ACSF treated slices were immediately grossly dissected to isolate 3-4 LHb tissues from both hemispheres and flash frozen in liquid nitrogen and stored at −80°C until further processing. All U50,488 treated LHb-containing slices were first incubated in ASCF + 10uM U50,488 at 28°C for 30min prior to gross-dissection and flash frozen. Samples were thawed, washed with PBS and sonicated in standard RIPA buffer containing protease and phosphatase inhibitors (Sigma). Supernatant was extracted following centrifugation and protein concentration was assessed using Pierce BCA Protein Assay Kit (Life Technologies). Samples with equal protein (20ug) content were combined with loading buffer and separated using standard gel electrophoresis on 4-20% polyacryllamide gel (Bio-Rad Techonolgies) and then partial-wet transferred to nitrocellulose membranes (BioRad). Total protein was first assessed using UV302nm activation (5min) prior to visualization at UV302nm for 30s (Azure Biosystems cSeries 280 Imager)[38]. Membranes were then rinsed and blocked with 5% BSA for 1hr prior to overnight incubation (4°C) with monoclonal rabbit anti-Phospho-p38 MAPK (Thr180/Tyr182) (D3F9)-XP (Cell signaling, 4511T, 1:1,000) or monoclonal rabbit anti-p38 MAPK (D13E1) XP (Cell signaling, 8690T, 1:1000). Membranes were then incubated with secondary antibody goat-anti rabbit IgG – HRP (Cell Signaling, 7074, 1:2,000), developed with Clarity Western ECL Substrate (Bio-Rad Laboratories) and detected using a BioRad ChemiDoc Touch image acquisition system (BioRad Laboratories) for 1-10min exposure time. All western blot analysis was performed using ImageJ and each sample was first normalized to its total protein concentration, and then normalized to the mean non-MD (control ACSF) and presented as fold-change from non-MD control. All data are presented as group mean ± SEM. For all qPCR data, non-MD and MD (PN21-28) LHb tissue was generated using the same methods used in WB and stored at −80C. Total RNA was isolated using TRIzol (Life Technologies)/choloroform-based extraction. Samples were quantified using NanoDrop spectrophotometry at 260 and 280 nm optical densities. Reverse transcriptase amplification of cDNA from total RNA was conducted with High-Capacity RNA-to-cDNA kit (Applied Biosystems) and Bio-Rad CFX96 Real-Time System. TaqMan gene expression assays (Thermo Fisher Scientific) with probes for *Oprk1* (cat. Rn0144892_m1) and actin (Actb, cat. Rn00667869_m1) as a control for total RNA and the comparative C_T_ (ΔΔC_T_) were used to assess *Oprk1* expression in relation to *Actb* control, as described in [13]. For each sample, the changes in target gene expression are given as 2^-ΔΔCT^. All data were normalized to the non-MD group and presented as fold change.

### 2.8. Statistics

Values are presented as means ± SEM. The threshold for significance was set at *P < 0.05 for all analyses. All statistical analyses of data were performed using Graphpad, Prism 8.4.1. For detecting the difference in distribution of silent, tonic or bursting LHb neurons in non-MD and MD rats, we used Chi-square tests. For neuronal excitability data (number of APs generated across current step) we used 2-way repeated measures (RM) ANOVA (RM in current step, and U50,488) of Ih- and Ih+ neurons separately. To assess whether all membrane and AP properties data (RMP, Rin, AP threshold, mAHP, fAHP, AP amplitude, and AP half-width) were normally distributed, we used the Shapiro-Wilk test (*α* =0.05) within each membrane/AP property across all data groupings (Ih and treatment). The effect of U50,488 on any AP/membrane properties were compared using Student’s paired-t-test, or nonparametric Wilcoxon paired test where appropriate. Baseline differences between Ih- and Ih+ groups were compared using unpaired Student’s t-test, or Mann-Whitney test (non-parametric data). All AP/membrane property group means and statistical test results are presented in table format: Supplemental Table 1(control non-MD: AP and intrinsic membrane properties with intact synaptic transmission), Supplemental Table 2 (control non-MD: AP and intrinsic membrane properties with blocked synaptic transmission), and Supplemental Table 3 (MD: AP and intrinsic membrane properties with intact synaptic transmission).

Mini Analysis software was used to detect and measure mIPSCs and mEPSCs using preset detection parameters of mIPSCs and mEPSCs with an amplitude cutoff of 5 pA. Effects of U50,488 on the mean and cumulative probabilities of mEPSC and mIPSC amplitude and frequency data sets were analyzed using paired student-t-tests and Kolmogorov-Smirnov tests(α=0.05), respectively.

The effects of MD on DynA (1-8) density were assessed using unpaired student t-test. All WB and qPCR data were analyzed first with Shapiro-Wilks test for normal distribution, followed by appropriate unpaired student t-test, or nonparametric Mann-Whitney U test.

## 3. Results and Discussion

### 3.1. KOR stimulation bidirectionally modulated LHb neuronal excitability

Compelling evidence implicates Dyn/KOR modulation of neuronal activity across a variety of reward/aversion processing brain regions [34, 45–48] that also project to the LHb. However, up until now the role of KOR signaling in the LHb has not been investigated. Here, we first sought to establish whether activation of KORs can regulate LHb neuronal activity and synaptic function within the LHb and then investigate how ELS may modulate this signaling within the LHb.

For selective KOR stimulation throughout this study, we used a KOR-specific agonist (U50,488, 10μM). First we tested the effects of U50,488 on LHb neuronal excitability by recording depolarization-induced AP generation in LHb neurons with intact synaptic transmission. We found that U50,488 had opposing effects on neuronal excitability in distinct LHb cell subpopulations that were discriminated by the presence or absence of Ih currents (i.e., Ih+ versus Ih-LHb neurons) **(Figure 1). Figure 1C** depicts the average current traces of Ih current recordings in response to a 50mV hyperpolarizing current step from LHb neurons of all control (non-MD) rats recorded throughout this study and a scatter plot of recorded Ih values across Ih- and Ih+ designations. U50,488 significantly decreased neuronal excitability in response to depolarization in Ih-LHb neurons as evident in the decreased average number of APs generated at each depolarizing current step compared to baseline levels **(Figure 1A).** Conversely, U50,488 significantly increased LHb neuronal excitability in Ih+ LHb neurons **(Figure 1B)**. Interestingly, we observed that the effects of U50,488 on LHb neuronal excitability persisted following a prolonged drug washout (data not shown). Although this may relate to a possible inability to completely wash the drug from the slice, it is also likely that acute KOR stimulation by U50,488 triggers a series of molecular adaptations in LHb neurons that induce and/or maintain a persistent change in neuronal excitability. To validate that U50,488 acted through KORs to change LHb neuronal excitability, we then repeated these experiments in the presence of a novel KOR-specific antagonist, BTRX-084266, 10-20μM (BlackThorn, Inc) which completely blocked the effects of the agonist, U50,488 **(Figure 1H-I)**.

**Figure 1.**
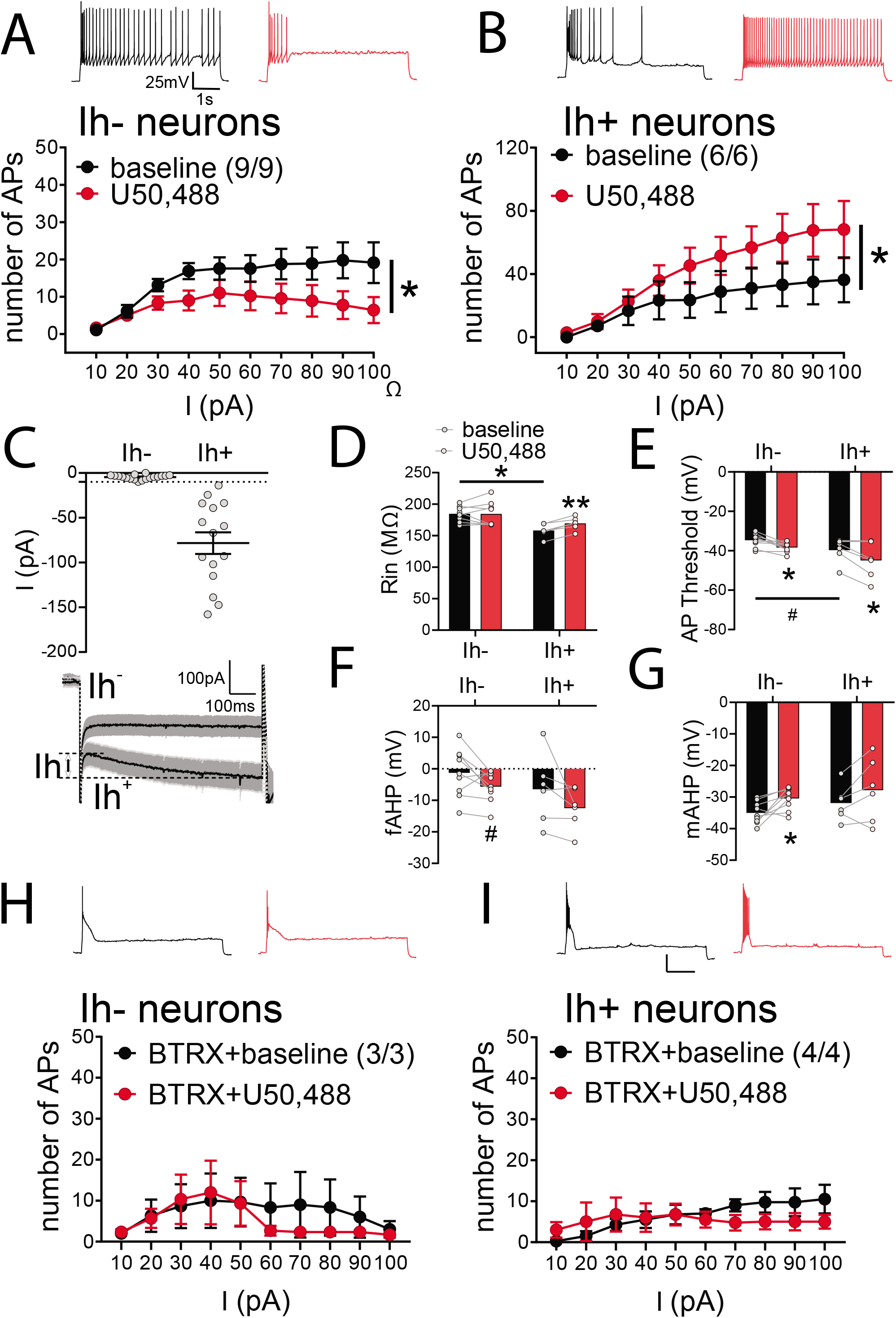
KOR agonist, U50,488, bidirectionally altered LHb neuronal excitability in distinct subpopulations of LHb neurons identified by Ih currents. **A-B**, inset: Example traces of AP generation in response to +50pA current injection at baseline (black) and post-U50,488 (red). **A.** U50,488 significantly decreased the number of AP generated across depolarizing current steps of Ih-neurons (2-way RM ANOVA, effect of U50,488: F(1,8)=7.3, p<0.0001; effect of current step: F(9,72)=5.6, p<0.05; U50,488xcurrent interaction: F(9,72)=2.6, p>0.05, n.s.) **B.** U50,488 significantly increased the number of AP generated across depolarizing current steps in Ih+ neurons (2-way RM ANOVA, effect of U50,488: F(1,5)=9.7, p<0.0001; effect of current step: F(9,45)=9.4, p<0.05; U50,488 x current interaction: F(9,45)=3.7, p<0.05) **C.** Scatter plot of Ih values across Ih-(n=19/14) and Ih+ (n=15/12) groupings; Mean ± SEM displayed as bar/whisker; traces: Ih current traces (mean [line] ± SEM [shaded region]) in response to a hyperpolarizing voltage step from −50mV to −100mV in Ih-(≤ −10pA, n=19/14) and Ih+ (> −10pA, n=15/12) groups. **D.** At baseline, Ih-neurons had larger input resistance (Rin) compared to Ih+ neurons (2-tailed unpaired t-test, t(12)=4.1, p<0.05) and U50,488 treatment significantly increased Rin only in Ih+ neurons (2-tailed paired t-test t(5)=7.3, p<0.001). **E.** At baseline, Ih-neurons had trending higher (less negative) AP thresholds compared to Ih+ neurons (2-tailed unpaired t-test, t(13)=2.0, p=0.062). U50,488 significantly lowered AP thresholds of both Ih-neurons (2-tailed paired t-test, t(8)=3.7, p<0.05) and Ih+ neurons (2-tailed paired t-test, t(5)=3.0, p<0.05). **F.** U50,488 induced a trending increase in fAHPs of Ih-neurons (2-tailed paired t-test, t(8)=1.9, p=0.095). **G.** U50,488 significantly decreased mAHPs of Ih-neurons (2-tailed paired t-test, t(8)=3.1, p<0.05). Graphs A-B, D-G: Ih-: n=9/9; Ih+: n=6/6. **H-I.** Pretreatment with a KOR-specific antagonist, BTRX-084266, prevented U50,488-induced changes in the excitability in Ih-neurons (n=3/3; 2-way RM ANOVA, effect of U50,488: F(1,2)=0.7, p>0.05 n.s.; effect of current: F(9,18)=2.0, p>0.05 n.s.; U50,488xcurrent interaction: F(9,18)=0.5, p>0.05 n.s.) and Ih+ neurons (n=4/4; 2-way RM ANOVA, effect of U50,488: F(1,3)=0.7, p>0.05 n.s.; effect of current: F(9,27)=0.3, p>0.05 n.s.; U50,488xcurrent interaction: F(9,27)=1.9 p>0.05 n.s.). All graphs: # p<0.1, * p<0.05, **p<0.001, ***p<0.0001. In this and subsequent figures n represents the number of recorded cells/rats.

In addition to measuring the number of APs in response to depolarization following KOR stimulation, we also assessed the effects of U50,488 on several intrinsic passive and active membrane properties across both Ih- and Ih+ LHb neurons derived from the recordings reported in Figure 1A-B **(Figure 1D-G and Supplemental Table 1)**. We found that at the baseline levels, Ih-neurons had significantly higher input resistance (Rin) than Ih+ neurons possibly due to the basal larger conductance of HCN/Ih channels in these neurons compared to Ih-neurons as Ih is active at resting membrane potentials in LHb neurons [49]. U50,488 only increased Rin of Ih+ neurons suggesting that KOR stimulation may result in possible decreases in resting K+ conductance that underlie membrane resistance such as leak K+ channels [50]. Their closure upon KOR stimulation could also promote the increased excitability of Ih+ LHb neurons, as seen in **Figure 1D**. Irrespective of opposing effects on the excitability of Ih- and Ih+ neurons, stimulation of KOR by U50,488 lowered AP thresholds in both Ih- and Ih+ neurons (**Figure 1E)**. Previously, we reported that another stress neuromodulator, corticotropin releasing factor (CRF), acts through CRFR1 to increase LHb intrinsic neuronal excitability uniformly across all LHb neurons [13]. This excitatory effect of CRF is also associated with increases in Rin and lower AP thresholds, similar to our findings in Ih+ LHb neurons following KOR activation. Part of the change in Rin and AP threshold by CRF-CRFR1 signaling is related to its effects on the small- (SK-) and large-(BK-) conductance K+ channels; which mediate mAHPs and fAHPs, respectively [13]. These K+ channels play an important role in regulating AP frequency of LHb neurons in response to depolarization. We previously observed that CRF-CRFR1 signaling increases fAHPs (while reducing mAHPs) in LHb neurons, thereby inducing a more hyperpolarized inter-spike membrane potential which could prevent voltage-gated Na+ channel inactivation and thus support the high frequency firing of LHb neurons during prolonged depolarization [13]. Here we found that while U50,488 did not affect fAHPs in either Ih- or Ih+ LHb neurons **(Figure 1F, Supplemental Table 1,** but also see **Figure 2E),** it significantly reduced mAHPs in Ih- LHb neurons with no effect on Ih+ LHb neurons **(Figure 1G)**. This may seem contradictory, as any reduction in mAHPs should promote rather than decrease neuronal excitability, but hyperpolarization induced by KOR stimulation as indicated by KOR-induced lowering of AP threshold potentials in Ih-neurons could also suppress SK channel activation and result in reduced mAHPs while still prevent the cell from reaching AP threshold and spiking. Altogether, these findings suggest that KOR activation may exert differential effects on LHb neuronal excitability through modulating numerous resting and active ion conductances that drive the membrane potential towards negative voltages. It is important to note that the changes induced by KOR on intrinsic measurements derived from the above AP recordings with intact synaptic transmission may not only be related to direct actions of KOR stimulation on LHb neurons, but also may involve KOR’s indirect effects. This could be related to KOR modulation of fast synaptic transmission (e.g., through GABA_A_Rs, AMPARs, NMDARs) as well as a complex interactions with various neuromodulators through their G-coupled protein receptors (GPCRs). Examples include CRF-CRFR1, GABA_B_Rs, metabotropic glutamate receptors (mGluRs), D2 and D4 DA receptors, and 5HT receptors which have all been shown to regulate intrinsic properties and intrinsic excitability of LHb neurons [13, 22, 24, 49, 51–56].

**Figure 2.**
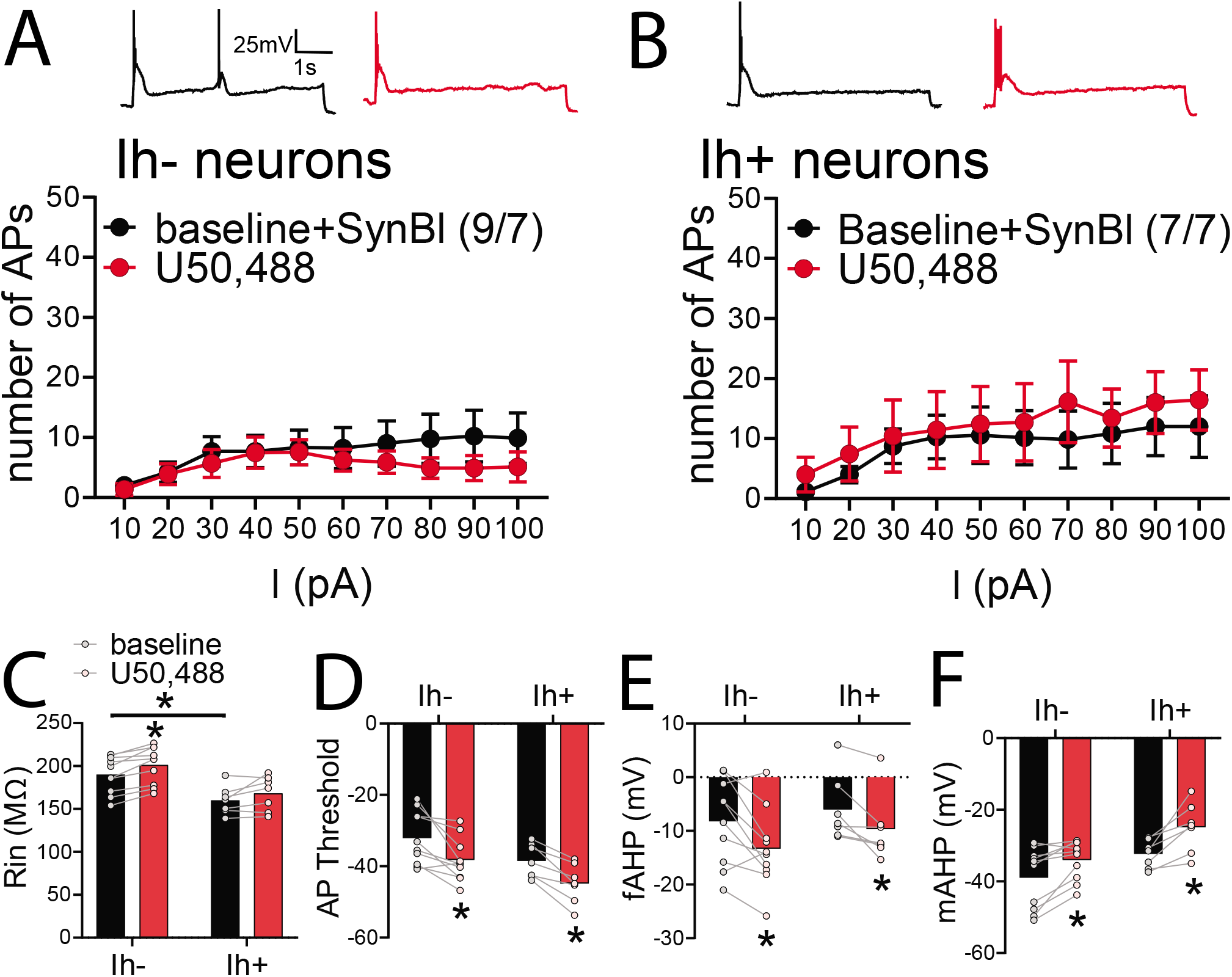
Effects of U50,488 on LHb neuronal excitability required intact synaptic transmission. **A-B.** inset: Example traces of AP generation in response to +50pA current injection at baseline (black) and post-U50,488 (red). U50,488 did not affect the number of AP generated across depolarizing current steps in **A.** Ih-neurons (2-way RM ANOVA, effect of U50,488: F(1,8)=1.7, p>0.05 n.s.; effect of current: F(9,72)=2.1, p<0.05; U50,488xcurrent interaction: F(9,72)=1.1, p>0.05 n.s.) or **B.** Ih+ neurons (2-way RM ANOVA, effect of U50,488: F(1,6)=0.5, p>0.05 n.s.; effect of current: F(9,54)=2.1 p<0.05; U50,488xcurrent interaction: F(9,54)=0.2, p>0.05 n.s.). **C.** At baseline, Ih-neurons had larger Rin compared to Ih+ neurons (2-tailed unpaired t-test, t(14)=3.0, p<0.05) and U50,488 significantly only increased Rin of Ih-neurons (2-tailed paired t-test, t(8)=5.0, p<0.05). **D.** U50,488 significantly decreased (more negative) AP thresholds in both Ih- (2-tailed paired t-test, t(8)=3.2 p<0.05) and Ih+ neurons (2-tailed paired t-test, t(6)=5.4, p<0.05). **E.** U50,488 significantly increased fAHPs of both Ih- (2-tailed paired t-test, t(8)=2.7, p<0.05) and Ih+ neurons (2-tailed paired Wilcoxon test, p<0.05). **F.** U50,488 significantly decreased mAHPs of both Ih- (2-tailed paired t-test, t(8)=3.0, p<0.05) and Ih+ neurons (2-tailed paired t-test, t(6)=4.0, p<0.05). All graphs: Ih-: n=9/7; Ih+: n=7/7, *p<0.05.

To our knowledge this is the first demonstration of functional KOR signaling in the LHb. Although the majority of LHb neurons is believed to be glutamatergic, there is more appreciation for anatomically and/or functionally diverse glutamatergic neuronal populations within the LHb that receive distinct synaptic inputs and have cell-specific projections to segregated and nonoverlapping targets including the VTA, substantia nigra compacta, rostromedial tegmental area (RMTg), or raphe nuclei (DR), thus comprising distinct neural output pathways from the LHb that control motivational behaviors [16, 19, 57–59]. Almost all LHb neurons projecting to the VTA or RN appear to express all four HCN subunits [60], however, a later study found that only a subset of glutamatergic medial LHb (mLHb) neurons with functional Ih are excited by DA-D4R activation of Ih currents and these neurons project to the RMTg but not to the VTA [49]. Interestingly, there is also an emerging and controversial notion of the existence of a small population of functional GABAergic interneurons within the LHb that locally suppress neuronal activity to increase motivation [61–63]. It possible that aversive or stressful stimuli can engage these distinct LHb neuronal populations through selective KOR-mediated activation of glutamatergic neurons and/ KOR-mediated inhibition of GABAergic neurons, ultimately increasing activity in glutamatergic output pathways, suppressing motivational behaviors, and promoting avoidance. Although whether the Ih- and Ih+ neurons identified in this study are glutamatergic or GABAergic remains to be directly assessed.

### 3.2 The effects of KOR stimulation on LHb neuronal excitability required intact fast synaptic transmission

Changes in both LHb neuronal intrinsic properties as well as synaptic transmission may drive the differential KOR modulation of LHb neuronal excitability in Ih- and Ih+ neurons. Therefore, we next tested whether KOR stimulation by U50,488 induces any change in intrinsic neuronal excitability in the absence of fast synaptic transmission by blocking AMPA, NMDA and GABA_A_ receptors. Under these conditions we observed that stimulation of KOR by U50,488 did not significantly affect neuronal excitability in response to depolarization in either Ih- or Ih+ LHb neurons **(Figure 2A-B).** Interestingly, under synaptic blockade conditions, some of our findings for KOR-mediated changes in intrinsic membrane properties persisted or were otherwise uncovered. For example, while we found that Ih-LHb neurons still exhibited a higher Rin compared to Ih+ LHb neurons, U50,488 now significantly increased the Rin of Ih-neurons. U50,488 also significantly lowered AP threshold, increased fAHPs and significantly decreased mAHP in both Ih- and Ih+ neurons **(Figure 2C-F, Supplemental Table 2)**. The persistence/appearance of effects of KOR-stimulation on intrinsic membrane properties without a significant effect on neuronal excitability highlights the necessity of presynaptic KOR stimulation on synaptic transmission rather than changes in intrinsic membrane properties of LHb neurons are required to drive KOR modulatory effects on LHb neuronal excitability.

### 3.3 KOR stimulation altered both glutamatergic and GABAergic synaptic transmission within the LHb

To directly test the effects of KOR signaling on synaptic transmission, we next tested the effects of U50,488 on AMPAR-mediated mEPSC **(Figure 3)** and GABAAR-mediated mIPSC **(Figure 4)**. It should be mentioned that recording of Ih is not feasible under mEPSC and mIPSC recordings given the necessary pharmacological manipulations and internal solutions required for these types of recordings also directly block or alter Ih currents. Therefore Ih- and Ih+ LHb neurons could not be distinguished in these recordings. We observed that U50,488 consistently (9 out of 10 neurons from 10 rats) reduced both the mean amplitude of mEPSCs as well as the cumulative probability of mEPSC amplitude (**Figure 3A**). Moreover, we found a significant reduction in the mean frequency of mEPSCs as well as a significant decrease in the cumulative probability of mEPSC inter-event interval (IEI) following U50,488 treatment (**Figure 3B)**. These findings indicate that KOR stimulation uniformly suppresses glutamatergic transmission within the LHb, significantly diminishing both presynaptic glutamate release and also decreasing postsynaptic AMPA receptor function (the number of AMPA receptors and/or the conductance of AMPA receptors) in LHb neurons.

**Figure 3.**
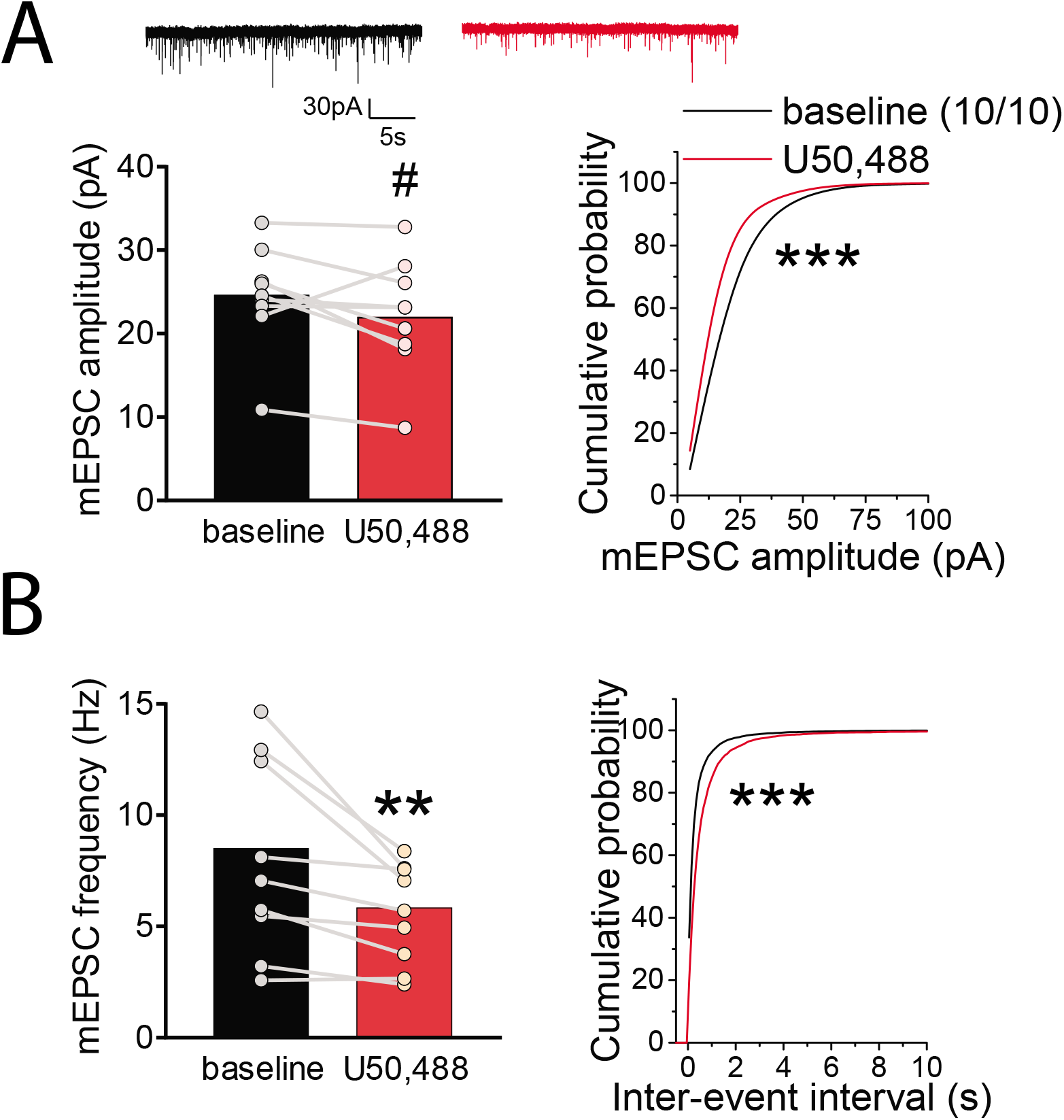
U50,488 suppressed glutamatergic transmission onto LHb neurons. **A.** Top: representative AMPAR-mediated mEPSC traces recorded from LHb neurons before (baseline, black) and after U50,488 (10μM, red) application. U50,488 had a trending decrease in the mean amplitude of mEPSCs (2-tailed paired-student t-test, t(9)=2.1 p=0.07) but significantly decreased cumulative probability of mEPSC amplitude (Kolmogorov-Smirnov [KS] test, p<0.0001). **B**. U50,488 treatment decreased the average mEPSC frequency (2-tailed paired-student t-test, t(9)=3.4, p<0.05) and cumulative probability of inter-event intervals of mEPSCs (KS test, p<0.0001). All data: n=10/10; #p<0.1, **p<0.01, ***p<0.0001.

The effects of KOR stimulation on GABAergic transmission was varied. While U50,488 significantly reduced the amplitude of mIPSCs, evident from the mean and cumulative probability plots of mIPSC amplitude (14/10 neurons/rats, **Figure 4A)**, its overall effects on the mean and cumulative probability of mIPSC frequency was either a decrease (10 out of the 14 neurons**, Figure 4B)** or an increase (4 out of the 14 neurons, **Figure 4C).** Given that that the effects of KOR stimulation on postsynaptic GABA_A_ receptor-mediated synaptic function was consistently diminishing across almost all LHb neurons similar to the effects we observed with postsynaptic AMPA receptor-mediated synaptic function, the variable effects of KOR signaling on presynaptic GABA release may suggest a projection-specific rather than a cell type-specific modulation of GABAergic synaptic transmission by presynaptic KOR signaling within the LHb.

**Figure 4.**
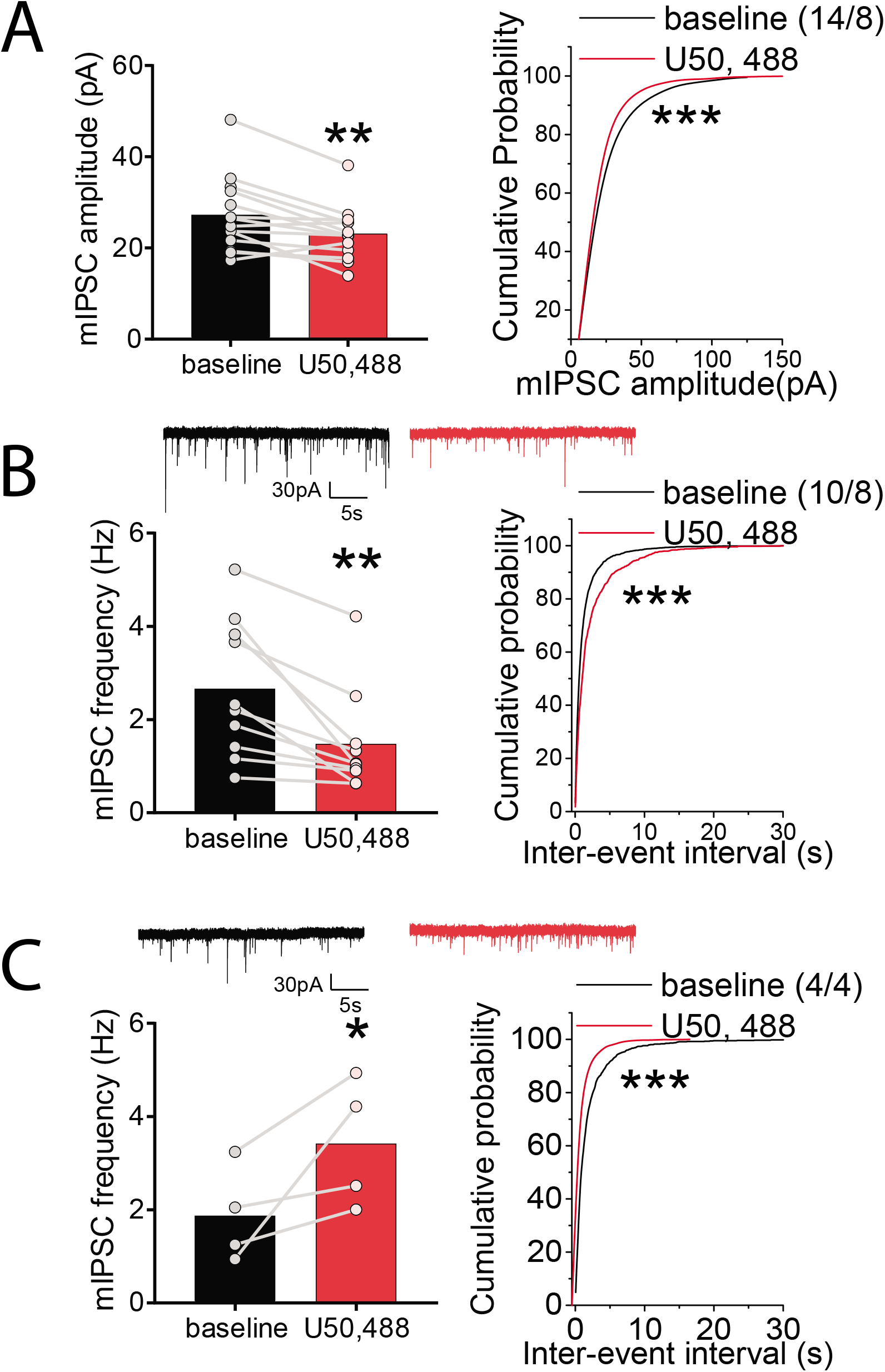
U50,488 altered GABAergic synaptic transmission onto LHb neurons. **A.** U50,488 decreased the mean amplitude of mIPSCs (n=14/8; 2-tailed paired-student t-test; t(13)=3.5, p<0.01) and decreased the cumulative probability of mIPSC amplitude (KS test, p<0.0001). **B**. Representative mIPSC traces at baseline (black) and following U50,488 treatment (red). Out of 14 recorded cells in **A**., U50,488 treatment decreased the average of mIPSC frequency (one-tail paired student t test, t(9)=3.9, p<0.01) as well as the cumulative probability of inter-event intervals of mIPSCs (KS test, p<0.0001)in 10 recorded cells (n=10/8). **C.** Representative mIPSC traces at baseline (black) and following U50,488 treatment (red). Out of 14 recorded cell in **A.**, U50,488 treatment increased the average frequency (one-tail paired student t test, t(3)=2.4, p<0.05) and the cumulative probability of inter-event intervals of mIPSCs (KS test, p<0.0001) in only 4 recorded cells (n=4/4). *p<0.05, **p<0.01, ***p<0.0001.

Our findings of KOR-induced suppression of presynaptic glutamatergic and GABAergic synaptic inputs to LHb neurons are consistent with previous studies of KOR activation in several brain regions including the hippocampus [64, 65], VTA [32, 33, 66–69], bed nucleus of the stria terminalis (BNST) [34, 70], NAc [45, 71], central amygdala neurons [72, 73] and ventrolateral periaqueductal gray [74]. Although general findings in the literature indicate that presynaptic KOR activation, through Gi/Go G-protein-coupled signaling, inhibits neurotransmitter release, our observation of KOR-induced increases in GABA release in a small subset of LHb neurons also raises the possibility of a cell- and/or pathway-specific modulation of GABA terminals by presynaptic KORs. Enhanced GABA release by KORs at specific inputs to the LHb, along with KOR induced-suppression of glutamate release, could contribute to the overall inhibitory effects of KOR signaling in suppressing Ih-LHb neuronal activity. Conversely, it is more likely that differential expression of KORs on various presynaptic inputs projecting selectively to Ih+ or Ih-LHb neurons biases excitatory/inhibitory drive (E/I balance) that may favor increased/decreased LHb neuronal excitability, respectively, as shown in our model (**Figure 8**).

In fact, this model is supported by studies in the NAc wherein activation of pre synaptic KORs decreases both excitatory and inhibitory synaptic drives onto NAc medium spiny neurons (MSNs), but the cell- and input-specific modification of EI balance in MSNs by KORs preferentially results in decreases in excitatory drive of D1 MSNs and disinhibition of D2 MSNs [45].

### 3.4. MD altered sensitivity of LHb neurons to KOR signaling

Previously we observed that MD blunts responses of LHb neurons to CRF signaling [11]. Given that dysregulation of Dyn/KOR modulation of DA circuits following stress including ELS is shown to mediate part of central CRF stress response or act in concert with CRF to promote drug-induced synaptic plasticity, drug seeking behaviors and stress-induced reinstatement [26–28, 48, 75, 76], we hypothesized that Dyn/KOR signaling within the LHb may also be dysregulated following MD stress. First, we recorded the spontaneous activity and firing patterns of LHb neurons in non-MD and MD rats in both cell-attached and whole cell recordings of APs to confirm our previous findings that MD significantly increases spontaneous LHb neuronal activity with more prevalence of active neurons in tonic and bursting modes of firing compared to non-MD controls in young-adolescent male rats (PN21-28) [7, 11] (**Figure 5).** We acknowledge that we could not measure Ih currents in all of our whole cell recordings for spontaneous LHb neuronal activity presented in Figure 5B. However, upon successful measurements of Ih in some of LHb neurons from non-MD and MD rats, we found an equal distribution of these currents across different modes of neuronal activity and firing patterns. Additionally, there was no significant difference in regard to the amplitude of Ih+ currents between Ih+ LHb neurons of non-MD and MD rats **(Figure 6C).** We then tested the effects of KOR stimulation by U50,488 on LHb neuronal excitability with intact synaptic transmission in MD rats. Strikingly, we found that MD completely abolished the effects of U50,488 on neuronal excitability in both Ih- and Ih+ neurons **(Figure 6A-B)**. Interestingly, the baseline difference of Rin between Ih- and Ih+ neurons that we observed with control non-MD rats was also present in MD rats and although U50,488 treatment did not affect Rin in either population, it significantly reduced mAHP of both Ih- and Ih+ LHb neurons in MD rats similar to non-MD controls (**Figure 6D-E, Supplemental Table 3)**. Considering that the effects of KORs on LHb neuronal excitability of control non-MD rats was absent when synaptic transmission was blocked **(Figure 2)**, the absence of these effects in MD rats suggests that the presynaptic effects of KOR stimulation on LHb neuronal excitability may be inhibited or occluded following MD. To directly address this, we next examined possible MD-induced alterations in Dyn/KOR signaling.

**Figure 5.**
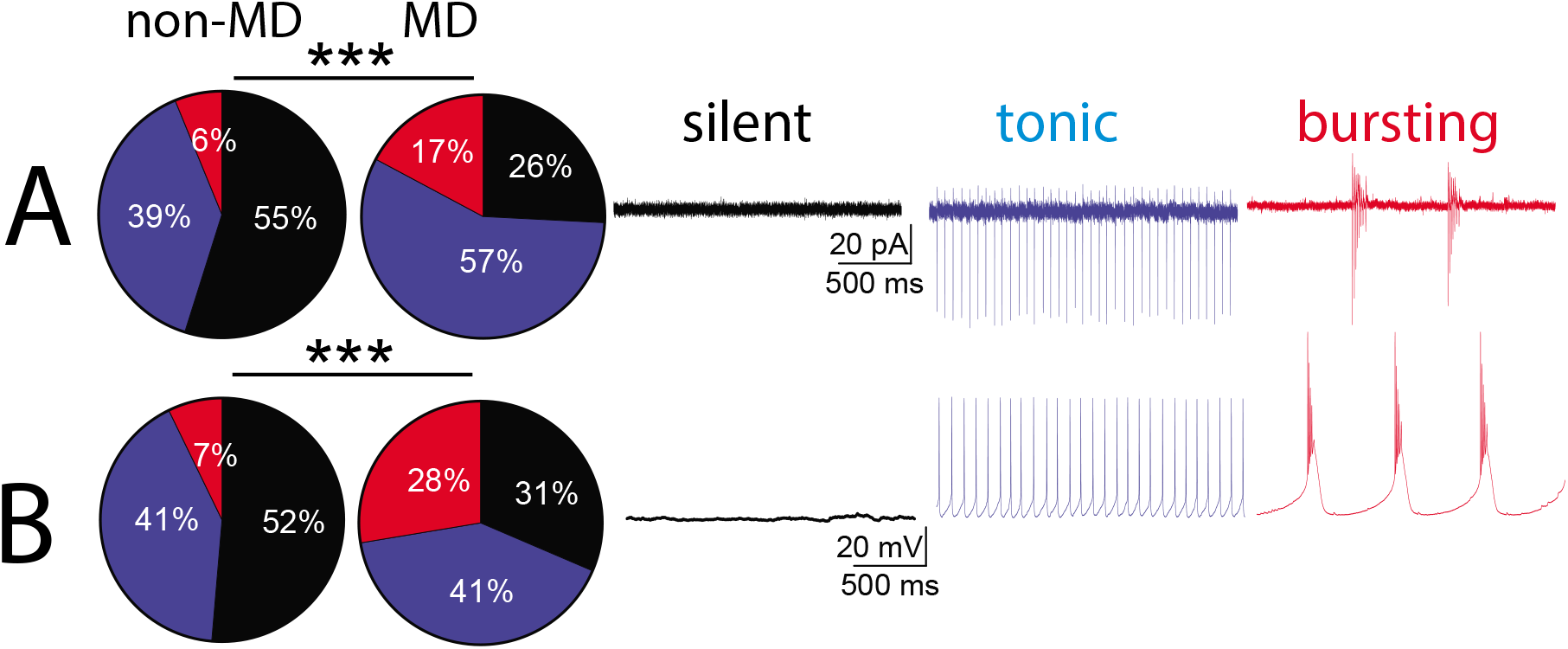
MD increased spontaneous activity of LHb neurons. **A-B**, Pie charts with representative traces of **A.** voltage-Clamp (VC, V=0) cell-attached recordings and **B.** current clamp (CC, I=0) whole-cell recordings of spontaneous neuronal activity across non-MD and MD rats. Comparison of the percent distributions of silent (black), tonic (blue), or bursting (red) LHb neurons showed a significant increase in tonic and bursting LHb neuronal activity following MD (Chi squared test, *** p<0.0001, VC: 113 neurons from 50 non-MD rats and 93 neurons from 48 MD rats; CC: 111 neurons from 48 non-MD rats and 105 neurons from 46 MD rats).

**Figure 6.**
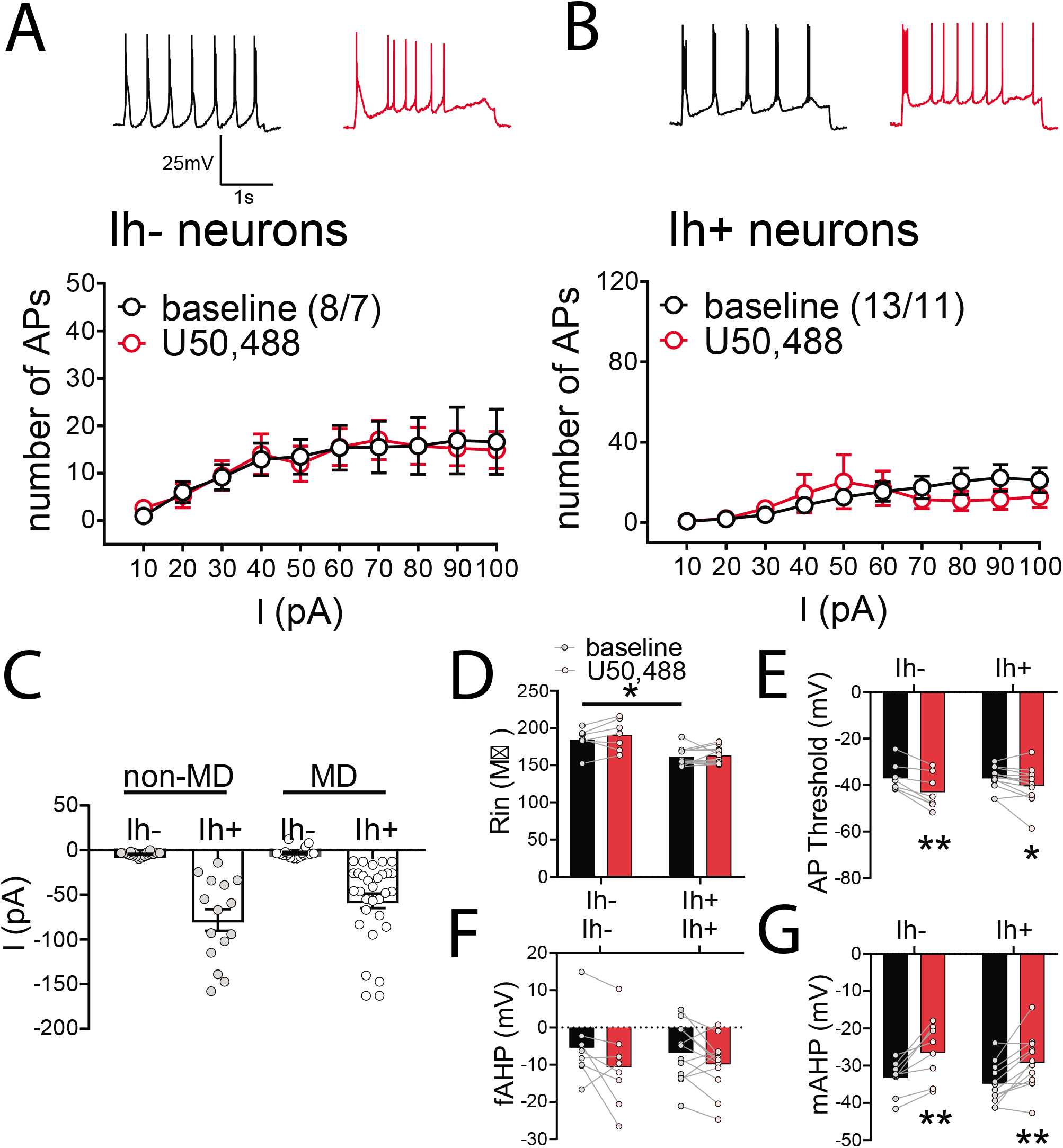
Effects of U50,488 on LHb neuronal excitability were absent following MD. **A-B.** Example traces of AP generation in response to +50pA current application at baseline (black) and 30min post-U50,488 (red). U50,488 did not significantly alter the number of AP generated across depolarizing current steps of **A.** Ih-neurons (2-way RM ANOVA, effect of U50,488: F(1,7)=0.04, p>0.05 n.s.; effect of current: F(9,63)=10.7, p<0.0001; U50,488xcurrent interaction: F(9,60)=0.3, p>0.05 n.s.) or **B.** Ih+ neurons (2-way RM ANOVA, effect of U50,488: F(1,11)=0.08, p>0.05 n.s.; effect of current: F(9,99)=3.7 p<0.0001; U50,488xcurrent interaction: F(9,99)=1.5, p>0.05 n.s.) in MD rats. **C.** Scatter plot of Ih currents recorded from LHb neurons across non-MD (grey) and MD (white) rats (non-MD, Ih-: n=19/14; MD, Ih-: n=8/7; non-MD, Ih+: n=15/12; MD, Ih+: n=12/10). There was no significant difference in the mean amplitude of Ih currents in Ih+ LHb neurons between non-MD and MD rats (Mann Whitney tests, p>0.05). **D.** At baseline, MD Ih-neurons had larger Rin compared to MD Ih+ neurons (2-tailed unpaired t-test, t(17)=3.6, p<0.05). **E.** U50,488 significantly lowered AP thresholds in both MD Ih-neurons (2-tailed Wilcoxon paired test, p<0.01) and MD Ih+ neurons (2-tailed paired t-test, t(11)=2.4 p<0.05). **F.** There was no effect of U50,488 on fAHPs in MD Ih-(2-tailed paired t-test, t(7)=1.9 p>0.05 n.s.) or MD Ih+ neurons (2-tailed paired t-test, t(11)=1.7, p>0.05 n.s.). **G.** U50,488 significantly decreased mAHPs of both MD Ih-neurons (2-tailed paired t-test, t(7)=4.3, p<0.01) and MD Ih+ neurons (2-tailed paired t-test, t(11)=4.1, p<0.01). All graphs (**A-B, D-G**: MD Ih-: n=8/7; MD Ih+: n=13/11), *p<0.05, **p<0.01, ***p<0.0001.

### 3.5 MD dysregulated DYN/KOR signaling within the LHb

The observed lack of LHb neuronal responses to KOR stimulation following MD may be due to changes in the expression of Dyn/KOR system or defective KOR downstream signaling. Therefore, we first examined whether MD alters the expression of Dyn-A (an endogenous opioid peptide that activates KORs with high potency) within the LHb. We performed double immunofluorescence with antibodies against NeuN (neuron-specific marker) and DYN-A(1-8) (an antibody specific to vesicular Dyn-A peptides, [77]) on male young-adolescent rats (PN28). Representative images of immuno-labeled DYN-A(1-8) highlight that the expression of this peptide was mainly seen within fibers across LHb **(Figure 7A).** We found that MD significantly and consistently increased DYN-A(1-8) density **(Figure 7A)** not only in PN28 but also in Juveniles (PN16) as well as adult (PN60) MD rats compared to non-MD (age-matched) indicating a persistent elevation in DYN expression across development **(Supplemental Figure 1)**.

**Figure 7.**
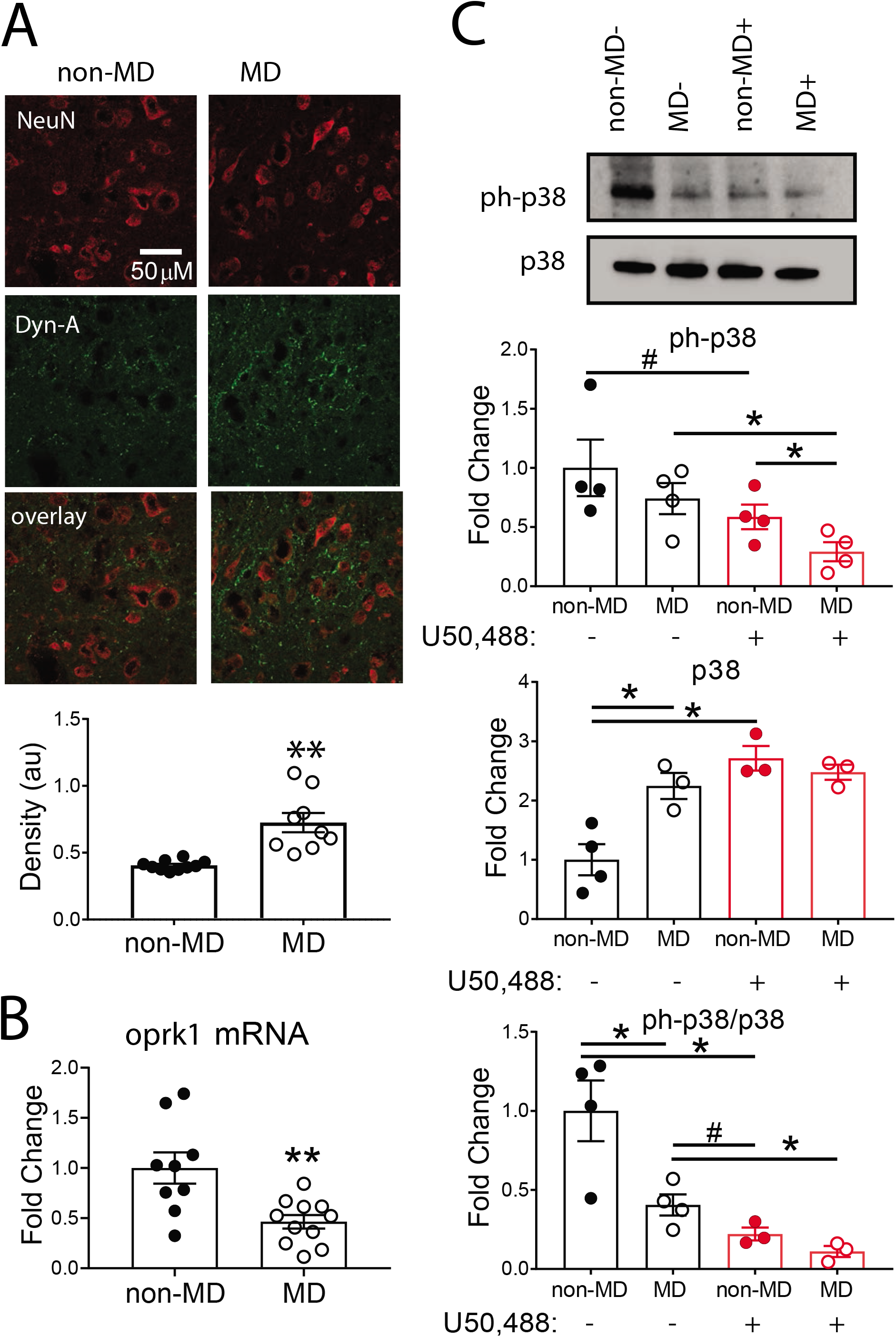
MD altered Dyn, KOR, and p38 (MAPK) expression in the LHb. **A.** Example images of non-MD (left) and MD (right) immunolabeling for LHb neurons (NeuN, red, top), Dyn-A(1-8) peptide (green, middle) and merged overlay images (bottom panel). Scale bar= 50μm. MD significantly increased average LHb Dyn-A density compared to non-MD controls (2-tailed unpaired t-test, t(17)=4.6, **p<0.001). **B.** LHb of MD rats had significantly lower abundance of opioid peptide receptor, kappa 1 *(oprk1)* mRNA expression compared to non-MD controls (unpaired student t-test, t(18)=3.4, p<0.01). **C.** inset: Example WB samples for ph-p38 (top) and total p38 (bottom) of LHb tissues across ACSF controls (-) and treatment with U50,488 (+) from non-MD/MD groups. **Top graph:** U50,488 significantly decreased ph-p38 in the LHb of MD rats (two-tailed unpaired t-test, t(6)=2.9, p<0.05) but only a trending effect was seen in the LHb of non-MD rats (one-tailed unpaired t-test, t(6)=1.6 p=0.081). ph-p38 levels in the LHb of MD+U50,488 group were significantly lower than non-MD+U50,488 group (one-tailed unpaired t-test, t(6)=2.2 p<0.05). All WB data: 3-4 rats each treatment. # p<0.1, * p<0.05, **p<0.01, *** p<0.0001.**Middle graph:** U50,488 significantly induced total p38 expression in the LHb of non-MD rats (one-tailed Mann Whitney test, p<0.05), while U50,488 incubation had no significant effect on the total p38 expression in the LHb of MD rats (two-tailed unpaired t-test, t(4)=0.9, p>0.05 n.s.). LHb of MD controls had significantly higher levels of total p38 expression compared to non-MD controls (two-tailed unpaired t-test, t(5)=3.4 p<0.05). **Bottom graph:** U50,488 significantly reduced ph-p38/p38 ratio in LHb of non-MD (two-tailed unpaired t-test, t(5)=3.4, p<0.05) and MD (two-tailed unpaired t-test, t(5)=3.5, p<0.05) rats. Additionally, LHb of MD rats had significantly lower Ph38/p38 compared non-MD controls (two-tailed, unpaired t-test t(6)=2.9, p<0.05). The ratio in LHb of MD+ACSF (MD controls) was trending higher than non-MD+U50,488 (2-tailed unpaired t-test, t(5)=2.1, p=0.086).

**Figure 8.**
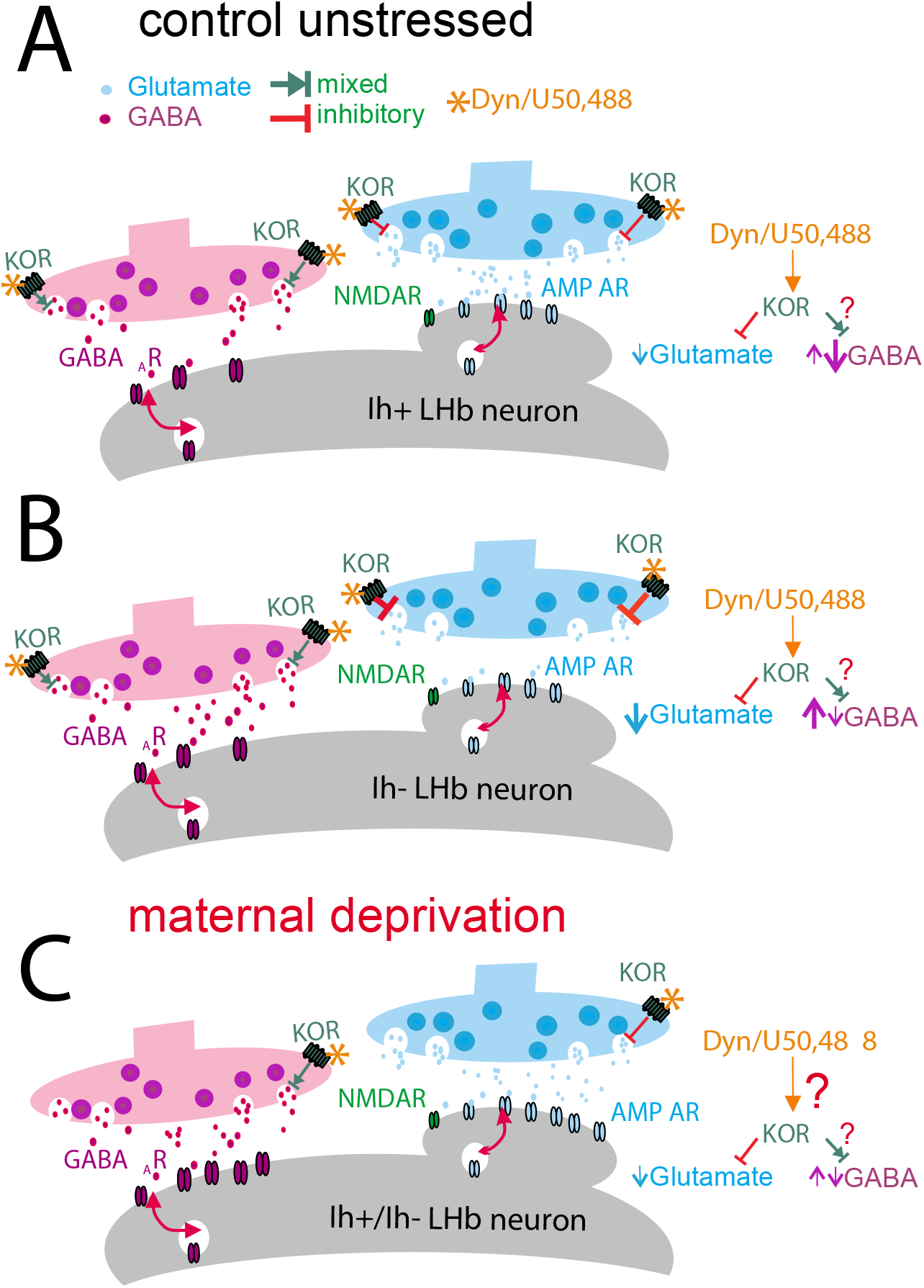
Dyn/KOR dysregulation of LHb neuronal activity following MD. The figure depicts our proposed model of Dyn/KOR differential regulation of LHb neuronal excitability in Ih+ and Ih-LHb neurons in control un-stressed rats (**A-B**) and its loss following MD (**C**). Dyn/KOR stimulation uniformly and presynaptically suppresses glutamatergic transmission (glutamate release) in LHb neurons while its effects on GABA release can be either a decrease (predominant effect) or even an increase (in small subset of neurons). We hypothesize that a preferential activation of KORs on GABAergic terminals and larger suppression of GABA release than glutamate release onto Ih+ LHb neurons results in KOR-induced increases in Ih+ Lhb neuronal excitability. In contrast, a preferential activation of KORs on glutamatergic terminal and greater suppression of glutamate release than GABA release in Ih-LHb neurons favors KOR-induced inhibition of Ih-Lhb neurons. It is also possible that in a subset of GABAergic terminals innervating Ih-neurons, KOR activation increases GABA release promoting inhibitory effects of KOR on Ih-LHb neurons. We speculate that MD stress engages Dyn/KOR signaling and upregulates this pathway with homeostatic decrease in KOR expression in response to Dyn hypertrophy which results in loss of proper responses of LHb neurons to this signaling.

Given MD-induced DYN signaling hypertrophy, it is possible that the lack of LHb neuronal responses to KOR stimulation of MD rats is driven by a downregulation of KORs in response to persistent activation of these receptors by hypertrophied Dyn transmission or alteration in signaling pathways down-stream of Dyn/KOR [25, 26, 32, 34, 45, 78]. In order to assess MD-induced alterations in the expression of KORs and/or KOR-specific downstream signaling, we measured KOR mRNA abundance and protein levels of the mammalian p38 mitogen-activated protein kinase (MAPK) and its phosphorylated (ph) active form (ph-p38), using qPCR and WB techniques, respectively. We found a significant decrease in *oprk1* (opioid peptide receptor kappa-1) mRNA expression in LHb tissues from MD rats compared to non-MD controls **(Figure 7B)**. This suggests that while Dyn-A expression is upregulated in the LHb by MD **(Figure 7A)**, there may be a homeostatic downregulation of receptor expression which could contribute to the blunted LHb neuronal responses to KOR activation following MD. Although it is important to note that mRNA abundance does not equate to protein expression or synaptic availability. While we found evidence that both Dyn-A peptide and KOR mRNA levels are altered following MD, it is likely that downstream molecular mechanisms from KORs were subject to change by MD. Several studies have shown that KOR activation by U50,488 (and Dyn) increases p38 MAPK signaling and can induce a transient increase in phosphorylation of p38 (ph-p38)[32, 78–80]. Therefore, as a proxy for KOR-activation, we tested whether MD and/or U50,488 bath application stimulated p38 signaling by measuring ph-p38 and total p38 protein levels using WB techniques. LHb tissues dissected from LHb slices from MD and non-MD rats were incubated in either ACSF (controls) or 10μM U50,488 (+) 30min prior to processing for WB analysis. Of note, while we chose the 30min incubation time to remain consistent with our electrophysiological data in this study, the transient increase in ph-p38 following U50,488 treatment generally peaks at 15min and by 30min the levels of ph-p38 are usually decreased [80]. Example blots for ph-p38 and p38 proteins are shown in **Figure 7C**, top inset. All ph-p38 and p38 density measurements were first normalized to the matching lane total protein signal before being averaged and normalized to fold change values relative to equivalent measures in LHb tissues from non-MD control rat samples. We thus compared the protein levels of ph-p38 and p38, as well as the ratio of ph-p38/p38, following U50,488 treatment in LHb tissues from non-MD and MD rats (**Figure 7C)**. We found that the levels of ph-p38 were similar in the LHb of non-MD control vs MD control rats. While ph-p38 was significantly decreased in MD+U50,488 compared to MD controls, there was a modest decrease with U50,488 treatment in the LHb of non-MD compared to non-MD controls that did not reach significance (**Figure 7C, bottom)**. In contrast to phosphorylation status, U50,488 significantly increased total p38 protein levels in the LHb of non-MD rats **(Figure 7C, middle)**. However, MD control rats exhibited higher baseline p38 total protein expression in the LHb compared to non-MD control rats, and U50,488 application did not further stimulate p38 expression in MD rats, indicating that decreased KOR expression may be a homeostatic response to higher basal activation of KORs by MD as evidenced by higher levels of Dyn-A and p38 expression in the LHb of MD rats by MD. Therefore, although the ratio of LHb ph-p38/p38 levels is significantly lower in MD control rats, the similar reduction in this ratio with U50,488 treatment in both MD and non-MD (**Figure 7C, top)** indicate that p38 signaling in the LHb of MD rats is still responsive to U50,488 KOR-induced kinase activation. Collectively, our molecular analysis of KOR/p38 signaling demonstrate that MD increases the total Dyn-A functional peptide expression in presynaptic terminal fibers in the LHb while downregulating KOR mRNA levels. Due to increased expression of p38 by MD, the sensitivity of the kinase to KOR stimulation to U50,488 remains intact. Therefore, MD appears to dysregulate Dyn/KOR signaling at multiple levels that involve projections from Dyn-containing brain areas to the LHb as well as the KOR expression/availability on synaptic terminals within the LHb, consistent with earlier studies of ELS [28, 76, 81–86].

## 4. Conclusion and future directions

To our knowledge our study is the first to demonstrate the existence of the functional KOR signaling in the LHb that differentially affects the excitability of discrete subpopulations of LHb neurons, mainly through changes in synaptic transmission. We also provided evidence for dysregulation of Dyn/KOR signaling following MD, used as a model of ELS. Given that KORs play an important role in drug- and stress-induced synaptic plasticity [26, 87, 88], our future studies will focus on KOR modulation of MD-induced intrinsic and synaptic plasticity within the LHb and at specific synaptic inputs to the LHb that may contribute to the later development of reward dysregulation and psychopathology associated with this model. We will also explore the therapeutic potential of targeting of KORs within the LHb in the context of ELS-induced neuropsychiatric and substance use disorders.

## Declarations of interest

none

## CRediT author statement

F.S.N, S.C.S and B.M.C designed the research; S.C.S, R.D.S, S.G, L.D.L performed the experiments. S.C.S, R.D.S, S.G, L.D.L and F.S.N analyzed the data and prepared the figures; S.C.S and F.S.N wrote the paper. All the authors contributed in reviewing and editing.

## Abbreviations

(AP): action potential,
(ACE): adverse childhood experiences,
(ACSF): artificial cerebral spinal fluid
(CRF): corticotropin-releasing factor,
(DA): dopamine,
(Dyn): dynorphin,
(ELS): early life stress,
(fAHP): fast-afterhyperpolarization current,
(HCN,Ih): hyperpolarization activated cation current,
(Rin): input resistance,
(IEI): interevent interval,
(KOR): Kappa opioid receptor,
(p38/MAPK): p38 mitogen-activated protein kinase,
(MD): maternal deprivation,
(mAHP): medium after hyperpolarization current,
(mEPSC): mini excitatory postsynaptic currents,
(mIPSC): mini inhibitory postsynaptic currents,
(non-MD): non-maternally deprived,
(NAc): nucleus accumbens,
(Ph-p38): phospho-p38,
(PN): postnatal age,
(RN): raphe nuclei,
(RMTg): rostromedial tegmental area,
(5HT): serotonin,
(VTA): ventral tegmental area

## Supplemental Tables and Figures

**Supplemental Figure 1. Effects of MD on Dyn-A expression in the LHb persisted across development.** Example images of Dynorphin A (Dyn-A 1-8) immunolabeling and quantification across ages **A**. PN16 and **B**. PN60 of non-MD and MD rats (PN16: n=6-7 rats/group, PN60: n=7-8 rats/group). For both, non-MD (left) and MD (right) immunolabeling for LHb neurons (NeuN, red, top), Dyn-A(1-8) peptide (green, middle) and merged overlay images (bottom panel) are shown. Scale bar= 50μm. MD significantly increased average LHb Dyn-A density compared to non-MD controls (PN16: two-tailed unpaired t-test, t(11)=2.04, *p<0.05; PN16: two-tailed unpaired t-test, t(12)=3.4, **p<0.01).

**Table 1. Effects of U50,488, on select membrane and AP properties on LHb neurons in slices from control non-MD rats in intact synaptic transmission.** Data represent measurements of resting membrane potential (RMP), AP threshold, input resistance (Rin), fast after-hyperpolarization current (fAHP), medium-after hyperpolarization current (mAHP), AP amplitude, AP half-width as combined (pooled data from Ih+ and Ih-neurons, left column), Ih-neurons (middle column) or Ih+ neurons (right column). line 1: mean±SEM, (neurons/rat); line 2: Result for passing (yes/no) the normal distribution test (Shapiro Wilks test, a=0.05); line 3 statistical summary: effect of U50,488 used paired student t-test (parametric) or Wilcoxon matched-pairs test (nonparametric) *p<0.05, **p<0.001, ***p<0.0001. Effect of Ih at baseline unpaired t-test (parametric) or Mann-Whitney U test (nonparametric) t<0.05, tt<0.001, ttt<0.0001; any trending effects denoted with # and p value stated.

**Table 2. Effects of U50,488 on membrane and AP properties of LHb neurons in slices from control non-MD rats with fast-synaptic transmission blocked.** Data are shown for similar measurements of intrinsic membrane and AP properties as presented in table 1 for combined (left column), Ih-neurons (middle column) or Ih+ neurons (left column). line 1: mean±SEM, (neurons/rat); line 2: Result for passing (yes/no) the normal distribution test (Shapiro Wilks test, a=0.05); line 3 statistical summary: (1) combined data: paired student t-test (parametric) or paired Mann-Whitney U test (nonparametric) (2) Across either Ih- or Ih+ data: mixed effects ANOVA (parametric) or RM Friedman’s test followed with paired student t-test or Mann Whitney U test, where appropriate. Effect of U50,488: *p<0.05, **p<0.001, ***p<0.0001, effect of Ih (baseline difference): t<0.05, tt<0.001, ttt<0.0001; Any trending effects denoted with # and p value stated.

**Table 3. Effects of U50,488 on membrane and AP properties of LHb neurons in slices from MD rats in intact synaptic transmission.** Data are shown for similar measurements of intrinsic membrane and AP properties as presented in table 1 for combined (left column), Ih-neurons (middle column) or Ih+ neurons (left column). line 1: mean±SEM, (neurons/rat); line 2: Result for passing (yes/no) the normal distribution test (Shapiro Wilks test, a=0.05); line 3 statistical summary: (1) combined data: paired student t-test (parametric) or paired Mann-Whitney U test (nonparametric) (2) Across either Ih- or Ih+ data: mixed effects ANOVA (parametric) or RM Friedman’s test followed with paired student t-test or Mann Whitney U test, where appropriate. Effect of U50,488: *p<0.05, **p<0.001, ***p<0.0001, effect of Ih (baseline difference): t<0.05, tt<0.001, ttt<0.0001; Any trending effects denoted with # and p value stated.

## Acknowledgements

The opinions and assertions contained herein are the private opinions of the authors and are not to be construed as official or reflecting the views of the Uniformed Services University of the Health Sciences or the Department of Defense or the Government of the United States. We are grateful to Drs. Irwin Lucki and Caroline Browne for their technical advice on qPCR experiments. The novel selective KOR antagonist (BTRX-084266) was generously provided by BlackThorn Therapeutics, Inc. We are especially thankful to Dr. Tanya Wallace from BlackThorn Therapeutics Inc. and Dr. Robert Koenig from the office of Technology Transfer at Henry M. Jackson Foundation for facilitating the Research Material Transfer of BTRX-084266.

## Funding

This work was supported by the National Institutes of Health (NIH)-National Institute of Drugs of Abuse (NIDA) Grant#R01 DA039533 to FSN. The funding agency did not contribute to writing this article or deciding to submit it.

## Notes

### Competing Interest Statement

The authors have declared no competing interest.

