## Supplemental Materials for "Early life stress dysregulates kappa opioid receptor signaling within the lateral habenula"

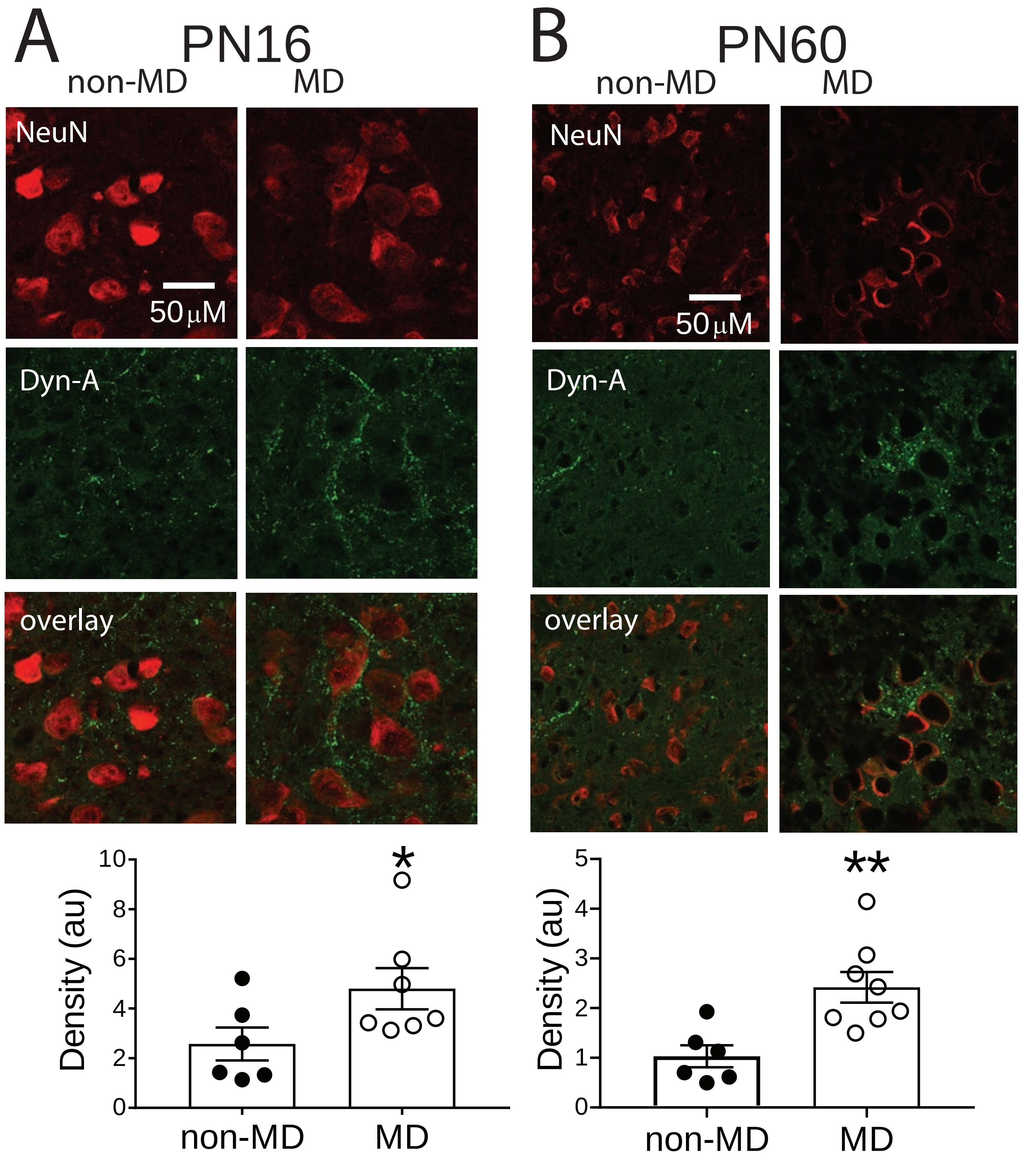


**Supplemental Figure 1:** DynorphinA (Dyn-A1-8) immunolabeling across ages A) PN16 and B) PN60 of non-MD and MD rats (PN16: n=6-7 rats/group, PN60: n=7-8 rats/group).

| | Combined | | Ih- | | Ih+ | |
|---|---|---|---|---|---|---|
| Property | baseline | U50,488 | baseline | U50,488 | baseline | U50,488 |
| Ih<br>parametric<br>significance | -20.0 ± 7.4, n14/12<br>no | NA | -3.4 ± 1.2, 9/9<br>yes<br>† | NA | -66.0 ± 19.1, 6/6<br>yes<br>† | NA |
| RMP<br>parametric<br>significance | -49.3 ± 2.9, n15/13<br>yes | NA | -52.2 ± 4.1, 9/9<br>yes | NA | -45.0 ± 3.3, 6/6<br>yes | NA |
| Rin<br>parametric<br>significance | 173.3 ± 4.7, n14/13<br>yes | 178.1 ± 4.7, n14/13<br>yes | 184.6 ± 4.4 n8/8<br>yes<br>† | 184.5 ± 7.0, n8/8<br>yes | 158.3 ± 4.4, n6/6<br>yes<br>† | 169.4 ± 4.1, n6/6<br>yes<br>** |
| AP Threshold<br>parametric<br>significance | -36.7 ± 1.3, n15/13<br>no | -41.0 ± 1.7, n15/13<br>no<br>** | -34.6 ± 1.2, n9/9<br>yes<br>†# p=0.0627 | -38.3 ± 0.8, n9/9<br>yes<br>* | -39.7 ± 2.5, n6/6<br>yes<br>†# p=0.0627 | -44.9 ± 3.7, n6/6<br>yes<br>* |
| fAHP<br>parametric<br>significance | -3.6 ± 2.4, n15/13<br>yes | -8.5 ± 1.7, n15/13<br>yes<br>* | -1.3 ± 2.6, n9/9<br>yes | -5.9 ± 1.8, n9/9<br>yes<br># p=0.0953 | -6.5 ± 4.5, n6/6<br>yes | -12.5 ± 2.7, n6/6<br>yes |
| mAHP<br>parametric<br>significance | -33.8 ± 1.2, n15/13<br>yes | -29.4 ± 1.7, n15/13<br>yes<br>** | -35.0 ± 1.2, n9/9<br>yes | -30.4 ± 1.2, n9/9<br>yes<br>* | -31.9 ± 2.3, n6/6<br>yes | -27.8 ± 4.1, n6/6<br>yes<br>* |
| AP amplitude<br>parametric<br>significance | 104.6 ± 3.0, n15/13<br>yes | 99.9 ± 2.9, n15/13<br>yes<br># p=0.0865 | 106.5 ± 3.4, n9/9<br>yes | 101.0 ± 3.1, n9/9<br>yes | 101.8 ± 5.8, n6/6<br>yes | 98.1 ± 5.7, n6/6<br>yes |
| AP halfwidth<br>parametric<br>significance | 1.9 ± 0.2, n15/13<br>yes | 1.8 ± 0.2, n15/13<br>yes | 2.2 ± 0.2, n9/9<br>yes<br>†# p=0.0687 | 2.1 ± 0.2, n9/9<br>yes | 1.6 ± 0.2, n6/6<br>yes<br>†# p=0.0687 | 1.3 ± 0.2, n6/6<br>yes |

Table 1. Membrane and AP properties of Ih- and Ih+ LHb neurons before and after U50,488 bath application in slices from non-MD control rats in intact synaptic transmission.

| | Combined | | Ih- | | Ih+ | |
|---|---|---|---|---|---|---|
| Property | basline | U50,488 | baseline | U50,488 | baseline | U50,488 |
| Ih | -14.6 ± 5.5, n15/12 | NA | -1.8 ± 1.3, n9/7 | NA | -33.9 ± 9.3, n6/6 | NA |
| parametric | no | | yes | | yes | |
| significance | | | †† | | †† | |
| RMP | -45.8 ± 2.9, n16/13 | NA | -44.1 ± 4.1 , n9/7 | | -47.9 ± 4.0 n7/7 | NA |
| parametric | yes | | yes | | yes | |
| significance | | | | | | |
| Rin | 177.2 ± 6.2, n16/13 | 186.4 ± 6.7, n16/13 | 190.5 ± 7.6, n9/7 | 200.6 ± 7.6, n9/7 | 160.1 ± 6.2, n7/7 | 168.2 ± 7.6, n7/7 |
| parametric | yes | yes | yes | yes | yes | yes |
| significance | | ** | † | ** | † | |
| AP Threshold | -35.3 ± 1.7, n16/13 | -41.6 ± 1.5, n16/13 | -32.8 ± 2.5, n9/7 | -39.2 ± 1.9, n9/7 | -38.5 ± 1.8, n7/7 | -44.8 ± 2.1, n7/7 |
| parametric | yes | yes | yes | yes | yes | yes |
| significance | | *** | | * | | * |
| fAHP | -6.5 ± 1.6, n16/13 | -11.0 ± 1.5, n16/13 | -6.8 ± 2.3, n9/7 | -11.9 ± 2.1, n9/7 | -6.0 ± 2.3, n7/7 | -9.7 ± 2.4, n7/7 |
| parametric | yes | yes | yes | yes | yes | no |
| significance | | ** | | * | | * |
| mAHP | -35.6 ± 1.9, n16/13 | -29.7 ± 1.9, n16/13 | -38.2 ± 2.9, n9/7 | -33.5 ± 1.9, n9/7 | -32.3 ± 1.5, n7/7 | -24.9 ± 2.6, n7/7 |
| parametric | no | yes | yes | yes | yes | yes |
| significance | | ** | | * | | ** |
| AP amplitude | 107.9 ± 1.5, n16/13 | 95.4 ± 1.6, n16/13 | 107.2 ± 1.8, n9/7 | 93.7 ± 2.6, n9/7 | 108.8 ± 2.7, n7/7 | 97.6 ± 1.6, n7/7 |
| parametric | yes | no | yes | yes | yes | yes |
| significance | | *** | | *** | | ** |
| AP halfwidth | 1.8 ± 0.1, n15/13 | 1.9 ± 0.1, n15/13 | 2.1 ± 0.1, n8/7 | 2.2 ± 0.1, n8/7 | 1.6 ± 0.2, n7/7 | 1.6 ± 0.2, n7/7 |
| parametric | yes | yes | yes | yes | yes | yes |
| significance | | | † | | † | |

Table 2. Membrane and AP properties of Ih- and Ih+ LHb neurons before and after U50,488 bath application in slices from non-MD control rats with fast synaptic transmission blocked.

| | combined | | Ih- | | Ih+ | |
|---|---|---|---|---|---|---|
| Property | baseline | U50,488 | baseline | U50,488 | baseline | U50,488 |
| Ih | -28.0 ± 5.9, n20/17 | NA | -4.1 ± 1.5, n8/7 | NA | -44.0 ± 6.4 n12/11 | NA |
| parametric | no | | yes | | yes | |
| significance | | | ††† | | ††† | |
| RMP | -47.6 ± 1.9, n20/17 | NA | -44.9 ± 2.4, n8/7 | NA | -49.5 ± 2.7 n12/11 | NA |
| parametric | yes | | yes | | yes | |
| significance | | | | | | |
| Rin | 169.7 ± 4.0, n19/17 | 173.3 ± 4.6, n19/17 | 184.1 ± 6.0, n7/7 | 190.7 ± 7.7, n7/7 | 161.3 ± 3.4, n12/11 | 163.1 ± 3.2, n12/11 |
| parametric | yes | no | yes | | yes | |
| significance | | | † | | † | |
| AP Threshold | -37.1 ± 1.2, n20/17 | -41.3 ± 1.7, n20/17 | -37.0 ± 2.2, n8/7 | -43.1 ± 2.6, n8/7 | -37.1 ± 1.4, n12/11 | -40.1 ± 2.2, n12/11 |
| parametric | yes | yes | no | yes | yes | yes |
| significance | | ** | | * | | * |
| fAHP | -6.2 ± 1.8, n20/17 | -10.2 ± 1.9, n20/17 | -5.4 ± 3.3, n8 | -10.6 ± 3.9, n8 | -6.8 ± 2.3, n12 | -9.9 ± 2.1, n12 |
| parametric | yes | yes | yes | yes | yes | yes |
| significance | | * | | | | |
| mAHP | -34.2 ± 1.1, n20/17 | -28.1 ± 1.6, n20/17 | -33.3 ± 1.7, n8 | -26.6 ± 2.6, n8 | -34.9 ± 1.5, n12 | -29.2 ± 2.1, n12 |
| parametric | yes | yes | yes | yes | yes | yes |
| significance | | *** | | * | | * |
| AP amplitude | 107.6 ± 1.4, n20/17 | 95.6 ± 2.6, n20/17 | 107.8 ± 2.3, n8 | 98.9 ± 4.6, n8 | 107.5 ± 1.7, n12 | 93.4 ± 3.0, n12 |
| parametric | yes | yes | yes | yes | yes | yes |
| significance | | *** | | # p=0.0591 | | ** |
| AP halfwidth | 1.9 ± 0.1, n20/17 | 2.0 ± 0.1, n20/17 | 2.1 ± 0.2, n8 | 2.3 ± 0.1, n8 | 1.7 ± 0.1, n12 | 1.8 ± 0.1, n12 |
| parametric | yes | yes | yes | yes | yes | yes |
| significance | | | †# p=0.0735 | | †# p=0.0735 | |

Table 3. Membrane and AP properties of Ih- and Ih+ LHb neurons before and after U50,488 bath application in slices from MD rats in intact synaptic transmission.
